# Stochastic effects during the establishment of β-lactam resistant *E. coli* mutants indicate conditions for collective resistance

**DOI:** 10.1101/2021.02.09.430436

**Authors:** Manja Saebelfeld, Suman G. Das, Arno Hagenbeek, Joachim Krug, J. Arjan G.M. de Visser

**Affiliations:** University of Cologne, Institute for Biological Physics, Cologne, Germany; Wageningen University, Laboratory of Genetics, Wageningen, The Netherlands

## Abstract

For antibiotic resistance to arise, new resistant mutants must establish in a bacterial population before they can spread via natural selection. Comprehending the stochastic factors that influence mutant establishment is crucial for a quantitative understanding of antibiotic resistance emergence. Here, we quantify the single-cell establishment probability of four *Escherichia coli* strains expressing β-lactamase alleles with different activity against the antibiotic cefotaxime, as a function of antibiotic concentration in both unstructured (liquid) and structured (agar) environments. We show that concentrations well below the minimum inhibitory concentration (MIC) can substantially hamper establishment, particularly for highly resistant mutants. While the pattern of establishment suppression is comparable in both tested environments, we find greater variability in establishment probability on agar. Using a simple branching model, we investigate possible sources of this stochasticity, including environment-dependent lineage variability. Lastly, we use the single-cell establishment probability to predict each strain’s MIC in the absence of social interactions. We observe substantially higher measured than predicted MIC values, particularly for highly resistant strains, which indicates cooperative effects among resistant cells at large cell numbers, such as in standard MIC assays.

## Introduction

A classical question in population genetics addresses the fate of a new adaptive allele in a population. Any new allele is initially present at a low frequency in the population and thus prone to extinction by genetic drift, even if the allele is beneficial (Patwa and Wahl 2008). In 1927, Haldane stated that a novel beneficial allele has to overcome drift loss and *establish* in the population before it can be picked up by selection; the probability of which he estimated to be roughly two times its selective benefit (Haldane 1927). Since then, Haldane’s approximation has been generalized and extended by a great number of models, but none of them has been tested empirically until about a decade ago (Patwa and Wahl 2008). With advances in genomic techniques, the issue of fixation probability has more recently been studied in experimental populations of microorganisms (e.g. Lang, Botstein, & Desai, 2011; Frenkel, Good, & Desai, 2014; Good, McDonald, Barrick, Lenski, & Desai, 2017). These studies looked at the fate of beneficial alleles once they reach a relatively high frequency in the population, finding that their fixation strongly depends on the competition with further beneficial alleles that appear via mutation over time. Only a small number of empirical studies has investigated the stochastic process of establishment of an initially rare beneficial allele arisen by *de novo* mutation in fungi (Gifford *et al*. 2013), bacteria (Farrell *et al*. 2017; Giometto *et al*. 2018) and nematodes (Chelo *et al*. 2013). Overall, these studies found that genetic drift dominates the establishment process, and the probability for a beneficial allele to establish in a population depends on the alleles’ initial frequency, its selective benefit, and the size of the population.

Whether a beneficial allele establishes successfully in a population is of particular relevance in cases of unwanted evolution, such as the evolution of antibiotic resistance. The worldwide growing concern about failing treatments of infections due to the emergence of resistant bacterial pathogens has led to the awareness that this problem must be tackled from different angles, including a better understanding of the emergence and evolution of antibiotic resistance (Baym, Stone, & Kishony, 2016; Smith et al., 2016; Furusawa, Horinouchi, & Maeda, 2018). Resistance alleles can be acquired via horizontal gene transfer or *de novo* mutation, resulting in the resistant allele initially being present in one or a few individuals within a susceptible population. But how likely is it that this single genotype can establish in the population and what factors influence this process? Using time-lapse microscopy, Coates et al. (2018) followed the growth of single *Escherichia coli* cells on agar, supplemented with different antibiotics. They found that the establishment of a single cell, determined by its growth into a visible macrocolony, was a highly stochastic process driven by fluctuations in cell death and birth, leading to the extinction of microcolonies at various time points. Similar results were found for two *Pseudomonas aeruginosa* strains, resistant to streptomycin or meropenem (Alexander and MacLean 2020). By seeding one to a few cells of a resistant strain in liquid medium with increasing antibiotic concentrations, it was found that at concentrations as low as 1/8 of the minimum inhibitory concentration (MIC) of this strain, its establishment probability was only 5% and that establishment of one cell was independent of the presence of other cells. The latter finding, however, may differ in systems where the antibiotic is broken down in the environment, leading to positive social interactions (Medaney *et al*. 2016).

A common resistance mechanism potentially allowing for social effects involves β-lactamase enzymes. β-lactamases hydrolyze the lactam ring of β-lactam antibiotics, which target penicillin-binding proteins (PBPs) that are involved in cell wall synthesis of gram-negative bacteria, thereby leading to loss of cell wall integrity and eventual cell lysis (Samaha-Kfoury & Araj, 2003; Bush, 2010). A well-studied and clinically relevant example is TEM-1 β-lactamase, which has high activity toward first-generation penicillins, but low activity toward third-generation β-lactams, including the cephalosporin cefotaxime (CTX). However, the TEM-1 allele is the ancestor of a large family of extended-spectrum β-lactamases, that have acquired mutations causing enhanced activity against more recently introduced penicillins and cephalosporins, including CTX (Salverda *et al*. 2010; Schenk *et al*. 2012; Van Dijk *et al*. 2017). As β-lactamases are expressed in the periplasmic space of gram-negative pathogens like *E. coli,* this system has been recognized as potentially cooperative, since the enzymatic breakdown in the periplasm reduces the antibiotic concentration in the environment (Brown *et al*. 2009), thereby cross-protecting susceptible bacteria nearby (Medaney *et al*. 2016). Such a cooperative behavior can be expected to have more pronounced consequences on agar due to the local breakdown of the antibiotic, creating microenvironments that may allow for the co-existence of strains with different levels of antibiotic resistance (Nicoloff & Andersson, 2016; Frost et al., 2018; Geyrhofer & Brenner, 2020).

Here, we examined the single-cell establishment probability of antibiotic-degrading bacteria under a range of antibiotic concentrations in both a structured (agar) and an unstructured (liquid) environment. We used four *E. coli* strains expressing β-lactamase alleles with different activities toward CTX, and hence different levels of resistance. Each strain was tested separately at low cell densities in both liquid and on agar medium with increasing CTX concentrations. To determine the single-cell establishment probability from the data, we developed a simple branching model, followed by a more detailed investigation of the cause of differences in stochasticity between the two environments. We show that environmental structure and cooperative behavior play at most a minor role in the establishment of resistant cells at low initial densities. Based on our results, we further introduce a new measure for resistance level that takes inoculum size and stochasticity into account. Application of the new measure to our data indicates substantial cooperative effects at higher cell densities, as typically used in standard MIC assays.

## Results and Discussion

In this study, we aimed at quantifying how the establishment of single bacterial cells with different levels of antibiotic resistance is affected by: (i) the antibiotic concentration, (ii) the environmental structure, and (iii) positive social interactions via the breakdown of the antibiotic in the environment. To determine single cell establishment probabilities, we performed one experiment in liquid (unstructured environment) and one on agar medium (structured environment). In each experiment, four *E. coli* strains (referred to as Ancestor, Single mutant, Double mutant, Triple mutant; cf. Table 1) with increasing levels of CTX resistance conferred by different antibiotic-hydrolyzing β-lactamase enzymes, were tested separately under exposure to a range of CTX concentrations. The tested concentrations differed between strains and were chosen to display establishment probabilities between 0 and 1 (see Material and Methods, and Supplement S1 - Table S1). We use the notation *p_e_*(*x*) to denote the probability that a single cell establishes a macroscopic population (i.e. turbidity in liquid or colony formation on agar, visible to the naked eye) at CTX concentration *x*. When the antibiotic concentration is zero, a population should establish with near certainty. Likewise, when the concentration is above the strains’ minimum inhibitory concentration (MIC), establishment is expected to be prevented. Stochastic outcomes can arise only at intermediate concentrations.

**Table 1:**
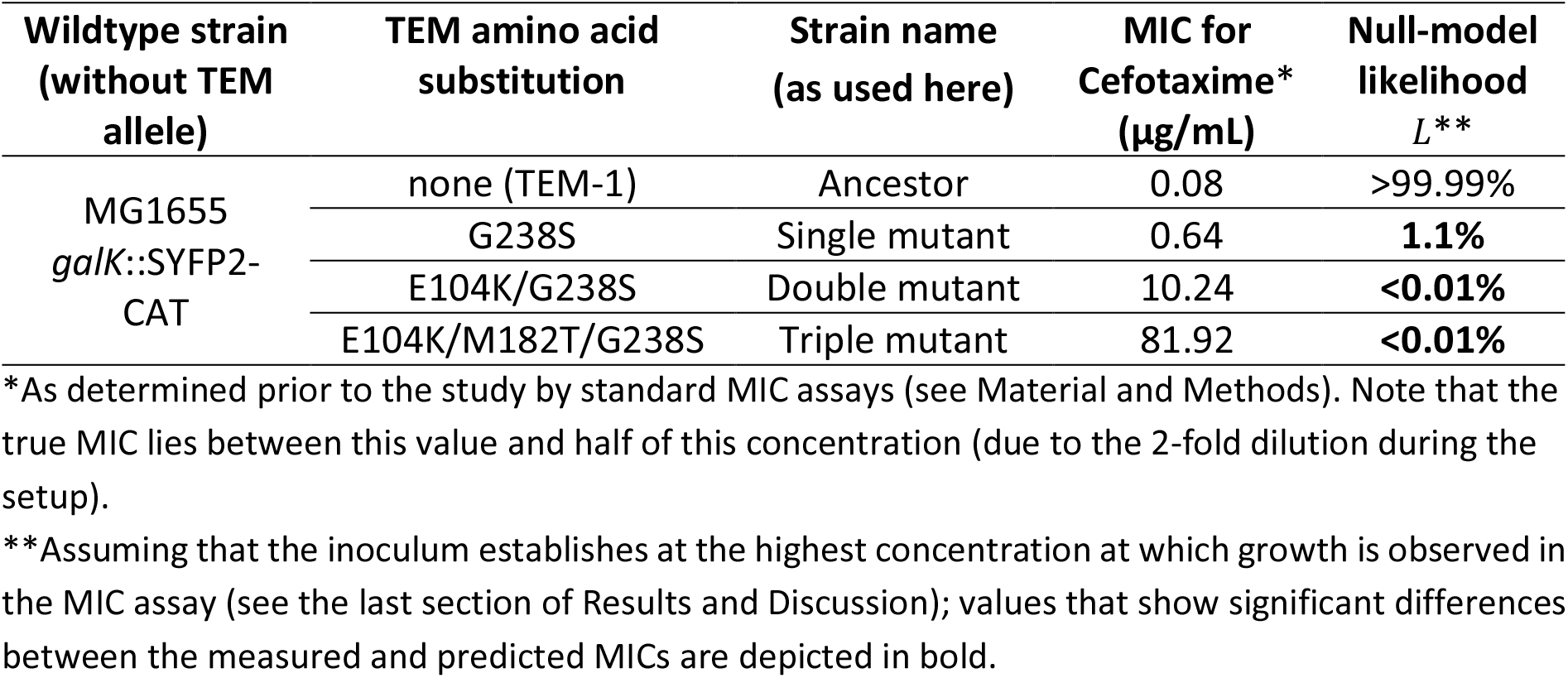
Overview of the used strains with their respective measured minimum inhibitory concentration (MIC) and null-model likelihood of the corresponding predicted MICs (see main text).

### Establishment probability declines well below the MIC in liquid

In the liquid medium experiment, we grew bacteria in multi-well plates and observed their establishment as a function of CTX concentration. Per strain, inoculum size (about 1 and 3 cells on average) and antibiotic concentration, a total of 285 replicate wells were tested, spread across three plates, including one medium control well per plate. Estimates of the viable cell numbers present in the inocula for evaluating the probabilities were obtained by applying cell culture aliquots on agar, taken from the culture dilutions that were used for inoculating the well plates. Using a bootstrapping algorithm and assuming a Poisson distribution of independently acting cells in each inoculum (details in Supplement S2), we estimated the single-cell establishment probability *p_e_*(*x*) for all strains and conditions. The results in Figure 1 show that the establishment probability nears zero well below each strain’s MIC and that this effect is more pronounced for strains with higher resistance levels.

**Figure 1.**
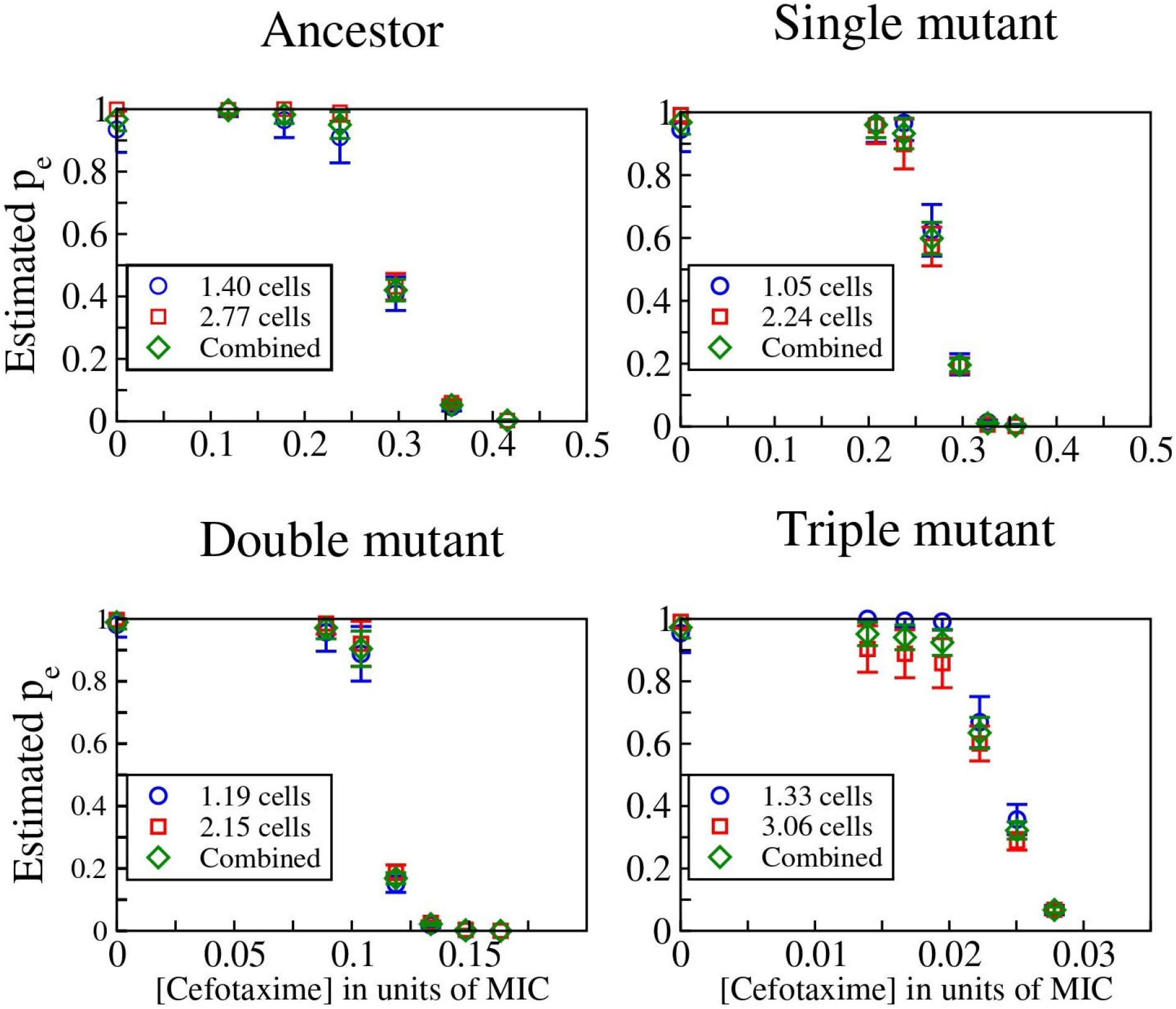
Single-cell establishment probabilities, *p_e_*, in liquid medium, estimated for the four tested strains as a function of antibiotic concentration, shown as a fraction of each strain’s measured MIC (cf. Table 1). For each strain, two inocula were used, one of approximately 1 cell (blue circles) and another of 2-3 cells (red squares); the average inoculum size estimated from separate CFU counts is shown. The inferred single-cell establishment probabilities are nearly identical for the two cases, implying that the establishment probability of a cell is not influenced by the presence of other cells at these low initial cell densities. The green diamonds are weighted averages over the two inoculum sizes (see Supplement S2 for details on the method used to estimate *p_e_*).

The observation that the establishment probability of a single cell nears zero at antibiotic concentrations well below the strain’s MIC has been shown before in *E. coli* for various bactericidal antibiotics (Coates *et al*. 2018) and in *P. aeruginosa* for streptomycin and meropenem (Alexander and MacLean 2020). These studies showed that the underlying causes for this are rooted in stochastic effects in population dynamics due to among-cell variability in death rate (Coates *et al*. 2018; Alexander and MacLean 2020) and lag-phase (Alexander and MacLean 2020), which become more pronounced with increasing antibiotic concentration. Therefore, the establishment probability is higher with increasing initial cell numbers, simply because more cells can survive and outgrow into a population (Coates *et al*. 2018; Alexander and MacLean 2020). This explains the decreased establishment probability of single cells at lower antibiotic concentrations compared to each strain’s MIC, as the latter is measured for inocula containing hundreds of thousands of cells per mL. While we confirm the results of this general pattern in our study, we extend the findings by using four *E. coli* strains with the same genetic background but different sensitivities toward the β-lactam antibiotic CTX. As these differences in sensitivity are based on differences in the level of β-lactamase activity (i.e. enzymatic capacity in breaking down CTX), our system provides the opportunity to study the effect of positive social interactions during establishment. Indeed, the increasingly pronounced discrepancy between the single-cell establishment probability and standard MIC estimates that is observed with increasing resistance level is, at least partially, attributed to an increased collective breakdown at high cell densities (see section “MIC and the inoculum effect” for details).

### Cells do not interact in liquid medium during establishment

One complication in estimating the single-cell establishment probability in liquid medium is that multiple cells may be present initially in some wells, and these cells may in principle interact, for example through competition for resources or the breakdown of antibiotic molecules. In that case, the establishment probabilities of individual cells would no longer be independent, biasing our estimates of *p_e_*(*x*). To test whether cells interact within initially small inocula, we conducted the liquid experiment with two different mean inoculum sizes (with approximately 1 and 3 cells, respectively) and estimated *p_e_*(*x*) for both cases under the assumption that there is no interaction (details in Supplement S2). If interactions occurred, the inferred values should disagree. However, Figure 1 shows that they agree closely, indicating that interactions between cells are insignificant at these low cell densities in unstructured environments.

The absence of social interactions in liquid medium at low cell density was also found in the study with *P. aeruginosa*, exposed to streptomycin or meropenem under similar conditions regarding culture volume and inoculum size (Alexander and MacLean 2020). In that system, positive social interactions are less likely to occur, because streptomycin resistance does not involve antibiotic-degrading enzymes. Our *E. coli* β-lactamase system on the contrary involves the potential for cooperative effects due to an active breakdown of the antibiotic in the environment. The absence of an effect in our data confirms that cell-to-cell interactions (both positive and negative) play an insignificant role during the establishment of single cells at such low densities in liquid medium and further shows that this observation is independent of the resistance mechanism.

### Establishment probability shows moderately higher stochasticity on agar

In the previous section, we found that the establishment probability of one cell was not influenced by the presence of other cells in liquid medium at very low densities. But how is this on agar? Previous studies have shown that evolutionary outcomes can largely differ depending on the environmental structure in the absence of migration (Perfeito *et al*. 2008; Lavrentovich *et al*. 2016; Santos-Lopez *et al*. 2019). For instance, while conditions in unstructured (liquid) environments commonly select for one strain to dominate the population, structured (agar) environments can maintain various strains due to the creation of microenvironments (Rainey and Travisano 1998; Kerr *et al*. 2002; Habets *et al*. 2007). One condition that can be affected locally on agar is the concentration of the antibiotic due to enzymatic breakdown. While it has been shown that this can result in cooperative effects, where a resistant strain protects more sensitive ones (Medaney *et al*. 2016; Rojo-Molinero *et al*. 2019) and such cooperation is more prominent in structured environments (Geyrhofer and Brenner 2020), it is, to the best of our knowledge, unknown whether such social interactions can influence the establishment of single antibiotic-resistant cells. To determine the effect of environmental structure on the establishment of β-lactamase-expressing cells, we plated about 200 cells of each *E. coli* strain on CTX-containing agar (eight plates per condition), followed by counts of colony-forming units (CFUs) after incubation. As for the experiment in liquid medium, the tested CTX concentration range depended on the strain to display establishment probabilities between 0 and 1 (see Material and Methods, and Supplement S1 - Table S1). The establishment probability was estimated as the CFU count relative to that at no CTX. The patterns of *p_e_* as a function of CTX concentration are broadly similar to those in liquid medium (Figure 2), indicating a minor role of environmental structure being involved in mutant establishment at low cell densities.

**Figure 2.**
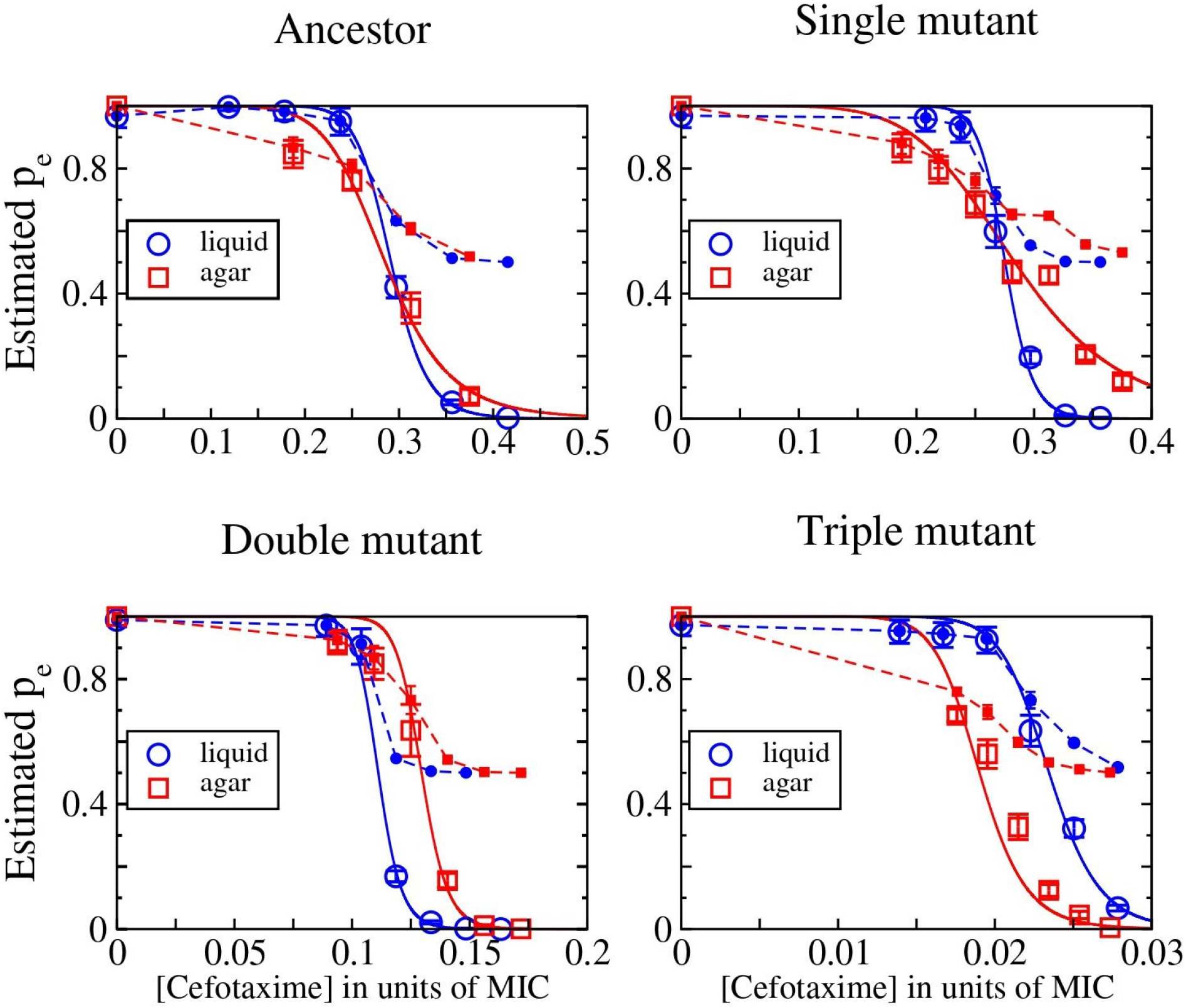
The single-cell establishment probability, *p_e_*, is shown as open symbols for the four tested strains in liquid (blue) and agar environments (red) for different CTX concentrations as the fraction of each strain’s measured MIC. For liquid medium, the combined results from both tested inoculum sizes (cf. Figure 1) are shown. The solid lines are fits performed with the Hill function *p_e_*(*x*) = (1 + (*x*/*x*_0_)*^n^*)^−1^ with x_0_ and *n* as parameters, where x_0_ is the concentration at which *p_e_* drops to 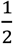, and *n* indicates the steepness of the curve. The values of the inferred parameters are given in Supplement S5, Table S2. The filled symbols connected by dashed lines show the cell division probability *p_d_* inferred from a simple branching model (see main text).

Although we found a general agreement of the establishment probability pattern between the liquid and agar experimental data, a closer comparison of the estimates of both environments shows that the decline of *p_e_* with CTX concentration is less steep on agar than in liquid (Figure 3a). This implies a higher degree of stochasticity on agar, since *p_e_* remains between 0 or 1 over a larger range of concentrations. We quantified this effect by calculating the Shannon entropy for the establishment probabilities in both environments and all four strains, finding that the entropy is higher on agar (Figure 3b).

**Figure 3.**
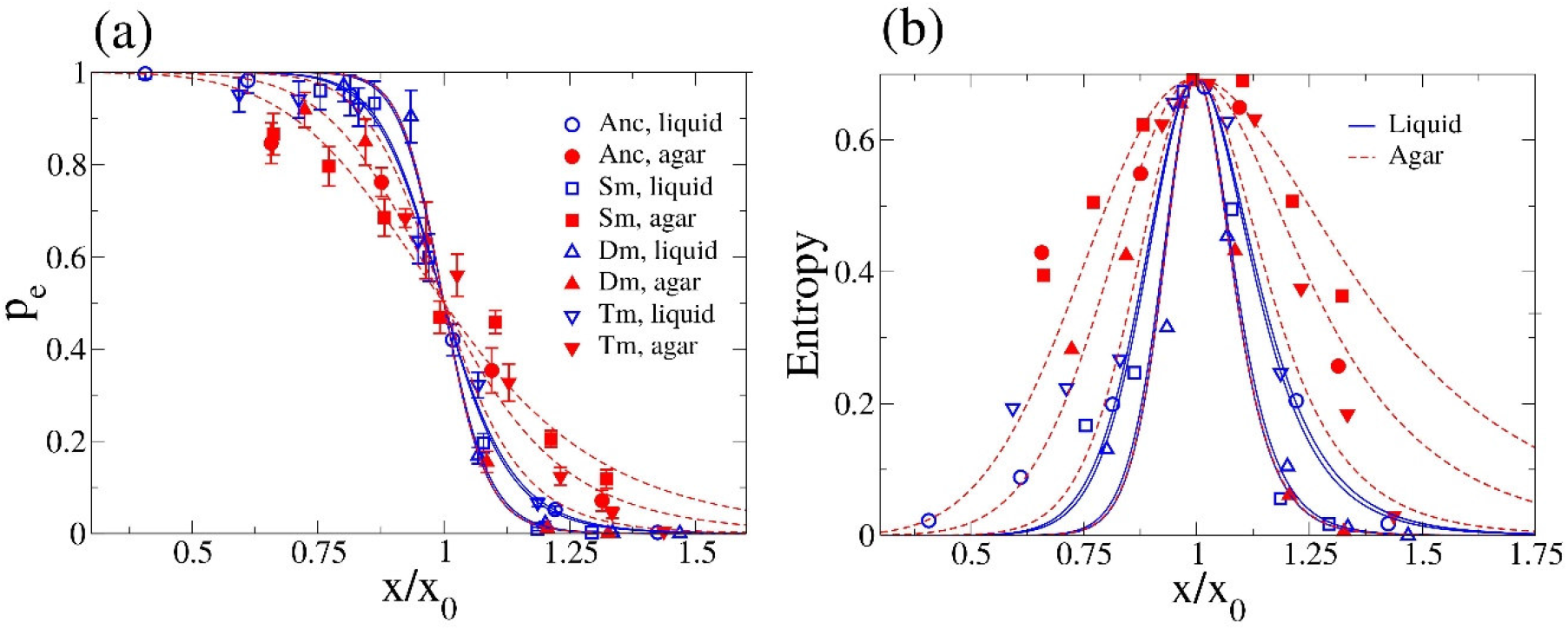
**(a)** Establishment probabilities for all the strains in both liquid and agar environments as a function of CTX concentration (Anc: Ancestor, Sm: Single mutant, Dm: Double mutant, Tm: Triple mutant), scaled by the corresponding estimate *x_0_* (i.e. the concentration at which *p_e_* drops to 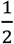). The curves for agar medium (red dotted lines) are less steep than those for liquid medium (blue solid lines). **(b)** The Shannon entropy is calculated and plotted for the *p_e_* values, using the standard expression *S* = –*p_e_* ln*p_e_* – (1 – *p_e_*)ln(1 – *p_e_*). It is seen to be higher for agar (red lines and symbols) compared to liquid (blue lines and symbols), as expected from the shallower slopes of the *p_e_*(*x*) curves for agar in panel (a).

Next, using a modelling approach we explored possible underlying mechanisms for stochasticity in the system. The simplest model is a branching process, where every cell divides with a fixed probability *p_d_*, or dies otherwise (see Supplement S3 for details), allowing us to estimate the division probability *p_d_* using the measured *p_e_* values. Figure 2 shows that under the simple branching model, the division probability appears to approach the value ½ asymptotically as opposed to the natural expectation of approaching zero asymptotically (see Figure S1 for an illustration of the possible shapes of the *p_e_* and *p_d_* curves in the simple branching model). This is a clear indication that the simple model does not fully capture the establishment process. One way to modify the model is to introduce variability in *p_d_* among cell lineages, and this fits the data reasonably well (see Supplement S4). Within this framework, the higher establishment stochasticity on agar could be due to small differences in *p_d_* of different cells on the agar surface (see Supplement S4 and Figure S3 for details). However, other possibilities remain; for instance, the cell division probability may change with time due to accumulating damages in cell wall structure under antibiotic action, changing rates of antibiotic and enzyme flux or changing CTX concentration over time at the location of a growing colony. Distinguishing conclusively between these alternatives is not possible based on our data, and we do not explore this issue any further here.

### MIC and the inoculum effect

Results from our experiments in liquid medium have shown a discrepancy between the CTX concentration where the single-cell establishment probability, *p_e_*, nears zero and the standard MIC value. As mentioned above, the reasons for this lie in the fact that standard MIC values are measured from large inocula of 5 × 10^5^ cells/mL, while establishment is a highly stochastic process at the cellular level. The stochastic nature of MIC estimates can be seen from the limited reproducibility of MIC assays, despite its coarse scale (Coates *et al*. 2018).

A conceptually cleaner approach to determine the MIC of a strain would therefore be to focus on a quantity that considers the role of stochasticity and initial inoculum size in the establishment of a population. For this purpose, we define the measure MIC*_q_*(*N*) as the minimum concentration at which an inoculum of initially *N* cells has a probability *q* of not being established (see Supplement S6). We suggest a null-model for this quantity, which assumes that the fates of the lineages arising from the N initial cells are independent, and therefore the extinction probability of the entire population equals (1–*p_e_*)*^N^*. Predictions based on this null-model can be used to test for the presence of interactions in the establishment process. For example, in a traditional MIC assay with an inoculum of 10^5^ cells/200μL, the null model predicts that 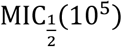 has the values of 0.053 (±3.5%), 0.31 (±4.7%), 2.0 (±3.6%), and 4.5 (±6.6%) μg/mL for the Ancestor, Single mutant, Double mutant and Triple mutant, respectively (see Supplement S6 for further details). The differences between these predicted values and the experimentally determined MICs (cf. Table 1) become increasingly pronounced with the level of resistance of the mutants. A choice of *q* other than ½ produces only modest changes in the predicted values (unless *q* is very close to the limits 0 or 1; see Supplement S6), and given the limited resolution of the MIC assay with two-fold increases in concentration, the precise choice of *q* does not affect the conclusion. The pattern of differences between the null-model 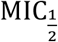 prediction and the measured MICs prompted us to calculate the likelihood, *L*, based on the null-model, that the inoculum establishes at the highest concentration at which growth is observed in the MIC assay (i.e. at half the reported MIC, due to the two-fold dilution steps). The results are shown in Table 1. While *L* is high for the Ancestor, it is low for all the mutants (*L* <2%), indicating that the measured MIC for the mutants is significantly above what we predict in the absence of social interactions (Table 1). This suggests that positive interactions among cells present at high density substantially enhance the collective establishment of mutant cell populations.

The effect of inoculum size on the MIC through social interactions has been recognized before (Brook 1989; Tan *et al*. 2012) and causes have been suggested to include the binding of the antibiotic to cellular components (Queenan *et al*. 2004) or the expression of proteins that inactivate the antibiotic (Lenhard and Bulman 2019). The four *E. coli* strains used here are genetically identical apart from the point mutations in the TEM allele, meaning that the number of penicillin-binding-proteins (PBPs), which are the main binding target of CTX (Piras *et al*. 1990; Cho *et al*. 2014), are comparable between them. Thus, binding of CTX molecules, although possibly contributing to the discrepancy between measured and predicted MICs, cannot explain the increasingly pronounced differences between the two resistance measures with increasing resistance levels of the mutant strains. As the level of resistance is conferred by increased enzyme activity against CTX with each additional point mutation (Knies *et al*. 2017), the collective breakdown of the antibiotic in the classical MIC assay is a likely explanation for the observed effect. Overall, our results on the discrepancy between the standard MIC value and the single-cell establishment probability are in line with the findings of other studies that criticized the MIC assay for dosage determination for the treatment of antibiotic-resistant infections (Gould & MacKenzie, 1997; Artemova et al., 2015; Alexander & MacLean, 2020).

## Conclusion

This study advances our understanding of the stochastic nature of the establishment of initially rare adaptive mutants (e.g. due to *de novo* mutation), a topic that has been under much theoretical scrutiny, but has rarely been investigated empirically. We find that the establishment of *E. coli* single mutant cells is negatively affected by CTX concentrations well below each of the four tested strain’s MIC, consistent with other recent findings (Alexander and MacLean 2020).

One aim of our study was to test if cooperative behavior via collective antibiotic breakdown among cells of the same strain, would affect the establishment probability and whether this would differ between unstructured and structured environments. Despite higher stochasticity on agar, our data show a reasonable agreement between the establishment in liquid and agar environments, indicating that environmental structure plays only a modest role, and that cooperative behavior has a negligible effect on the establishment probability of single resistant mutant cells in isolation. However, the process of establishment of antibiotic resistant mutants is particularly relevant for *de novo* mutants arising in populations of relatively susceptible bacteria, and in recent work, we demonstrated positive net effects on mutant establishment from susceptible cells in the same experimental system (Saebelfeld et al. 2021).

Lastly, we point out limitations of the traditional MIC measurements that make MIC values highly sensitive to the assay conditions. We suggest a more robust measure of resistance, MIC*_q_*(*N*), which includes both inoculum size and probabilistic effects, for which we constructed a null model based on the single-cell establishment probability, *p_e_*, by assuming homogeneous cell populations and no social interactions across lineages. The null-model prediction disagrees with the measured MIC in systematic ways, indicating strong cooperative effects at high cell densities (Saebelfeld *et al*. 2021). We propose that this measure may be used as a reference for detecting conditions of collective resistance for other bacterial strains and antibiotics.

## Material and Methods

### Strains and culture conditions

Four strains were used for the experiments, derived from *Escherichia coli* strain MG1655 *galK*::SYFP2-CAT (a kind gift from the lab of Dan Andersson via Peter Lind). This strain (DA28100) had previously been modified with a YFP (yellow fluorescent protein) marker cassette, containing a resistance gene for chloramphenicol (Gullberg *et al*. 2014). For the current study, the chloramphenicol resistance was removed, and each one of four TEM variants was inserted into the *galK* locus, together with the pTac promoter from plasmid pACTEM (Barlow and Hall 2002): the ancestral TEM-1 allele (referred to as “Ancestor”), and three mutant alleles with either 1, 2 or 3 point mutations in the TEM allele (referred to as “Single mutant”, “Double mutant” and “Triple mutant”, respectively). While the ancestral TEM-1 allele confers only low activity towards the β-lactam antibiotic cefotaxime (CTX), the three mutants show increasing CTX resistance with each additional substitution (Table 1). All TEM loci are under the control of the LacI repressor and are expressed by adding 50 μM Isopropyl β-D-1-thiogalactopyranoside (IPTG) to the growth medium. The minimum inhibitory concentration (MIC) of each strain for CTX had previously been determined in duplicates (Table 1), using 2-fold increases in CTX concentration in microtiter plates filled with 200 μL M9 minimal medium (containing 0.4% glucose, 0.2% casaminoacids, 2 μg/mL uracil and 1 μg/mL thiamine) and 50 μM IPTG, inoculated with 10^5^ cells and incubated for 24h at 37°C.

For culturing, all strains were first recovered from glycerol stocks by streaking them out on LB (Luria-Bertani) agar plates and incubated overnight at 37°C. From those plates, one colony was picked and introduced into 1 mL M9 medium (as above) and incubated overnight at 37°C, 250 rpm. The cultures were then serially diluted with phosphate-buffered saline (PBS) to the density needed for the particular experiment (see below), assuming an initial density of 2.5×10^9^ cells/mL.

### Liquid experiment

Each of the four strains was tested separately in the absence of CTX and six CTX concentrations (Supplement S1, Table S1), where the particular tested concentrations depended on the strains’ resistance level. The range of concentrations was chosen to display establishment probabilities between 0 and 1, based on pilot experiments to find the right conditions (data not shown).

Per strain and CTX concentration, on average either about 1 or 2-3 cells were seeded into 285 wells across three 96-well microtiter plates. For this, 190 μL M9 medium with the respective cefotaxime concentration and 50 μM IPTG were pipetted into all wells. Overnight cultures of the strains were diluted to 200 and 100 cells/mL with PBS. 10 μL of the dilutions were pipetted into the wells. One well per plate served as medium control, mock-inoculated with 10 μL PBS. The plates were incubated for about 40 h at 37°C (static). After incubation, OD at 600 nm of all wells was measured in a plate reader (Victor3^™^, PerkinElmer) without the plate lid. All medium controls across the experiment showed no sign of growth; the OD ranged from 0.033 to 0.038. Thus, a threshold of >0.05 was applied to determine whether a population successfully established.

To determine the mean number of seeded cells in culture dilutions, aliquots of the dilutions were dropped on LB agar plates (120 × 120 mm). Per dilution and strain, a total of 3×32 10 μL drops were applied to the plates, let dry for about 10 to 20 minutes, incubated at 37°C until leaving, and then moved to 30°C overnight to prevent overgrowth, followed by counting the number of colonies per droplet.

### Agar experiment

As in the liquid experiment, all strains were tested separately and the CTX concentration range depended on the strain. Five to eight concentrations were tested per strain (Supplement S1, Table S1). The overnight cultures were diluted to 4000 cells/mL. 50 μL (~200 cells) of the dilutions were spread onto 92 mm agar plates containing M9 medium (as above) with 1.5 % agar, 50 μM IPTG and the respective cefotaxime concentration, using a bacterial spreader. Per strain and cefotaxime treatment, eight replicate plates were used. The plates were incubated at 37°C until colonies were big enough to count them unambiguously (20 to 48 hours). Each colony is regarded as a successfully established single cell.

## Acknowledgements

We greatly thank Andrew Farr for providing the fluorescently-labelled MG1655 strain (which were a generous gift from the Dan Andersson lab) with the removed chloramphenicol resistance as well as the protocol for inserting the TEM alleles; Helen Alexander for fruitful discussions of her work, advice on the liquid experiment setup and valuable input on an earlier version of the manuscript; and Tobias Bollenbach for advice on the *p_e_* model.

## References

Alexander H. K., and R. C. MacLean, 2020 Stochastic bacterial population dynamics restrict the establishment of antibiotic resistance from single cells. Proc. Natl. Acad. Sci. U. S. A. 117: 19455–19464. https://doi.org/10.1073/pnas.1919672117

Artemova T., Y. Gerardin, C. Dudley, N. M. Vega, and J. Gore, 2015 Isolated cell behavior drives the evolution of antibiotic resistance. Mol. Syst. Biol. 11: 822. https://doi.org/10.15252/msb.20145888

Barlow M., and B. G. Hall, 2002 Predicting evolutionary potential: *in vitro* evolution accurately reproduces natural evolution of the TEM β-lactamase. Genetics 161: 1355.

Baym M., L. K. Stone, and R. Kishony, 2016 Multidrug evolutionary strategies to reverse antibiotic resistance. Science (80−.). 351. https://doi.org/10.1126/science.aad3292

Brook I., 1989 Inoculum Effect. Rev. Infect. Dis. 11: 361–368. https://doi.org/10.1093/clinids/11.3.361

Brown S. P., S. A. West, S. P. Diggle, and A. S. Griffin, 2009 Social evolution in micro-organisms and a Trojan horse approach to medical intervention strategies. Philos. Trans. R. Soc. B Biol. Sci. 364: 3157–3168. https://doi.org/10.1098/rstb.2009.0055

Bush K., 2010 Bench-to-bedside review : the role of β-lactamases in antibiotic-resistant Gram-negative infections. Crit. Care 14: 224.

Chelo I. M., J. Nédli, I. Gordo, and H. Teotónio, 2013 An experimental test on the probability of extinction of new genetic variants. Nat. Commun. 4: 1–8. https://doi.org/10.1038/ncomms3417

Cho H., T. Uehara, and T. G. Bernhardt, 2014 Beta-lactam antibiotics induce a lethal malfunctioning of the bacterial cell wall synthesis machinery. Cell 159: 1300–1311.

Coates J., B. R. Park, D. Le, E. Şimşek, W. Chaudhry, et al., 2018 Antibiotic-induced population fluctuations and stochastic clearance of bacteria. Elife 7: 1–26. https://doi.org/10.7554/eLife.32976

Dijk T. Van, S. Hwang, J. Krug, J. A. G. M. De Visser, and M. P. Zwart, 2017 Mutation supply and the repeatability of selection for antibiotic resistance. Phys. Biol. 14. https://doi.org/10.1088/1478-3975/aa7f36

Farrell F. D., M. Gralka, O. Hallatschek, and B. Waclaw, 2017 Mechanical interactions in bacterial colonies and the surfing probability of beneficial mutations. J. R. Soc. Interface 14. https://doi.org/10.1098/rsif.2017.0073

Frenkel E. M., B. H. Good, and M. M. Desai, 2014 The fates of mutant lineages and the distribution of fitness effects of beneficial mutations in laboratory budding yeast populations. Genetics 196: 1217–1226. https://doi.org/10.1534/genetics.113.160069

Frost I., W. P. J. Smith, S. Mitri, A. San Millan, Y. Davit, et al., 2018 Cooperation, competition and antibiotic resistance in bacterial colonies. ISME J. 12: 1582–1593. https://doi.org/10.1038/s41396-018-0090-4

Furusawa C., T. Horinouchi, and T. Maeda, 2018 Toward prediction and control of antibiotic-resistance evolution. Curr. Opin. Biotechnol. 54: 45–49. https://doi.org/10.1016/j.copbio.2018.01.026

Geyrhofer L., and N. Brenner, 2020 Coexistence and cooperation in structured habitats. BMC Ecol. 20: 1–15. https://doi.org/10.1186/s12898-020-00281-y

Gifford D. R., J. A. G. M. De Visser, and L. M. Wahl, 2013 Model and test in a fungus of the probability that beneficial mutations survive drift. Biol. Lett. 9. https://doi.org/10.1098/rsbl.2012.0310

Giometto A., D. R. Nelson, and A. W. Murray, 2018 Physical interactions reduce the power of natural selection in growing yeast colonies. Proc. Natl. Acad. Sci. U. S. A. 115: 11448–11453. https://doi.org/10.1073/pnas.1809587115

Good B. H., M. J. McDonald, J. E. Barrick, R. E. Lenski, and M. M. Desai, 2017 The dynamics of molecular evolution over 60,000 generations. Nature 551: 45–50. https://doi.org/10.1038/nature24287

Gould I. M., and F. M. MacKenzie, 1997 The response of *Enterobacteriaceae* to beta-lactam antibiotics-’round forms, filaments and the root of all evil’. J. Antimicrob. Chemother. 40: 495–499.

Gullberg E., L. M. Albrecht, C. Karlsson, L. Sandegren, and D. I. Andersson, 2014 Selection of a multidrug resistance plasmid by sublethal levels of antibiotics and heavy metals. MBio 5: 19–23. https://doi.org/10.1128/mBio.01918-14.Editor

Habets M. G., T. Czaran, R. F. Hoekstra, and J. A. G. de Visser, 2007 Spatial structure inhibits the rate of invasion of beneficial mutations in asexual populations. Proc. R. Soc. B Biol. Sci. 274: 2139–2143.

Haldane J. B. S., 1927 A mathematical theory of natural and artificial selection, Part V: selection and mutation. Math. Proc. Cambridge Philos. Soc. 23: 838–844. https://doi.org/10.1017/S0305004100015644

Kerr B., M. A. Riley, M. W. Feldman, and B. J. M. Bohannan, 2002 Local dispersal promotes biodiversity in a real-life game of rock-paper-scissors. Nature 418: 171–174. https://doi.org/10.1038/nature00823

Knies J. L., F. Cai, and D. M. Weinreich, 2017 Enzyme efficiency but not thermostability drives cefotaxime resistance evolution in TEM-1 β-lactamase. Mol. Biol. Evol. 34: 1040–1054. https://doi.org/10.1093/molbev/msx053

Lang G. I., D. Botstein, and M. M. Desai, 2011 Genetic variation and the fate of beneficial mutations in asexual populations. Genetics 188: 647–661. https://doi.org/10.1534/genetics.111.128942

Lavrentovich M. O., M. E. Wahl, D. R. Nelson, and A. W. Murray, 2016 Spatially constrained growth enhances conversional meltdown. Biophys. J. 110: 2800–2808. https://doi.org/10.1016/j.bpj.2016.05.024

Lenhard J. R., and Z. P. Bulman, 2019 Inoculum effect of β-lactam antibiotics. J. Antimicrob. Chemother. 74: 2825–2843. https://doi.org/10.1093/jac/dkz226

Medaney F., T. Dimitriu, R. J. Ellis, and B. Raymond, 2016 Live to cheat another day: bacterial dormancy facilitates the social exploitation of β-lactamases. ISME J. 10: 778–787. https://doi.org/10.1038/ismej.2015.154

Nicoloff H., and D. I. Andersson, 2016 Indirect resistance to several classes of antibiotics in cocultures with resistant bacteria expressing antibiotic-modifying or -degrading enzymes. J. Antimicrob. Chemother. 71: 100–110. https://doi.org/10.1093/jac/dkv312

Patwa Z., and L. M. Wahl, 2008 The fixation probability of beneficial mutations. J. R. Soc. Interface 5: 1279–1289. https://doi.org/10.1098/rsif.2008.0248

Perfeito L., M. I. Pereira, P. R. A. Campos, and I. Gordo, 2008 The effect of spatial structure on adaptation in *Escherichia coli*. Biol. Lett. 4: 57–59. https://doi.org/10.1098/rsbl.2007.0481

Piras G., A. El Kharroubi, J. Van Beeumen, E. Coeme, J. Coyette, et al., 1990 Characterization of an *Enterococcus hirae* penicillin-binding protein 3 with low penicillin affinity. J. Bacteriol. 172: 6856–6862. https://doi.org/10.1128/jb.172.12.6856-6862.1990

Queenan A. M., B. Foleno, C. Gownley, E. Wira, and K. Bush, 2004 Effects of inoculum and beta-lactamase activity in AmpC- and extended-spectrum beta-lactamase (ESBL)-producing *Escherichia coli* and *Klebsiella pneumoniae* clinical isolates tested by using NCCLS ESBL methodology. J. Clin. Microbiol. 42: 269–275. https://doi.org/10.1128/JCM.42.1.269

Rainey P. B., and M. Travisano, 1998 Adaptive radiation in a heterogeneous environment. Nature 394: 69–72.

Rojo-Molinero E., M. D. MacIà, and A. Oliver, 2019 Social behavior of antibiotic resistant mutants within *Pseudomonas aeruginosa* biofilm communities. Front. Microbiol. 10: 1–11. https://doi.org/10.3389/fmicb.2019.00570

Saebelfeld M., S. G. Das, J. Brink, A. Hagenbeek, J. Krug, et al., 2021 Antibiotic breakdown by susceptible bacteria enhances the establishment of β-lactam resistant mutants. Front. Microbiol. 12. https://doi.org/10.3389/fmicb.2021.698970

Salverda M. L. M., J. A. G. M. de Visser, and M. Barlow, 2010 Natural evolution of TEM-1 β-lactamase: experimental reconstruction and clinical relevance. FEMS Microbiol. Rev. 34: 1015–1036. https://doi.org/10.1111/j.1574-6976.2010.00222.x

Samaha-Kfoury J. N., and G. F. Araj, 2003 Recent developments in β lactamases and extended spectrum β lactamases. Bmj 327: 1209–1213. https://doi.org/10.1136/bmj.327.7425.1209

Santos-Lopez A., C. W. Marshall, M. R. Scribner, D. J. Snyder, and V. S. Cooper, 2019 Evolutionary pathways to antibiotic resistance are dependent upon environmental structure and bacterial lifestyle. Elife 8: 1–23. https://doi.org/10.7554/elife.47612

Schenk M. F., I. G. Szendro, J. Krug, and J. A. G. M. de Visser, 2012 Quantifying the adaptive potential of an antibiotic resistance enzyme. PLoS Genet. 8: e1002783. https://doi.org/10.1371/journal.pgen.1002783

Smith E., C. A. Lichten, J. Taylor, C. MacLure, L. Lepetit, et al., 2016 Evaluation of the EC action plan against the rising threats from antimicrobial resistance.

Tan C., R. Phillip Smith, J. K. Srimani, K. A. Riccione, S. Prasada, et al., 2012 The inoculum effect and band-pass bacterial response to periodic antibiotic treatment. Mol. Syst. Biol. 8: 1–11. https://doi.org/10.1038/msb.2012.49

