## Supplementary material for "Stochastic effects during the establishment of β-lactam resistant *E. coli* mutants indicate conditions for collective resistance"

### S1. Experimental conditions

**Table S1:** Cefotaxime concentrations used per strain in the liquid and agar experiments.

| | **Strain*** | **Cefotaxime concentration (µg/mL)** | | | | | | | |
|---|---|---|---|---|---|---|---|---|---|
| *Liquid experiment* | **Ancestor** | 0 | 0.01 | 0.015 | 0.02 | 0.025 | 0.03 | 0.035 | - |
| | **Single mutant** | 0 | 0.14 | 0.16 | 0.18 | 0.20 | 0.22 | 0.24 | - |
| | **Double mutant** | 0 | 0.96 | 1.12 | 1.28 | 1.44 | 1.6 | 1.76 | - |
| | **Triple mutant** | 0 | 1.20 | 1.44 | 1.68 | 1.92 | 2.16 | 2.40 | - |
| *Agar experiment* | **Ancestor** | 0 | 0.015 | 0.02 | 0.025 | 0.03 | - | - | - |
| | **Single mutant** | 0 | 0.12 | 0.14 | 0.16 | 0.18 | 0.20 | 0.22 | 0.24 |
| | **Double mutant** | 0 | 0.96 | 1.12 | 1.28 | 1.44 | 1.6 | 1.76 | - |
| | **Triple mutant** | 0 | 1.44 | 1.60 | 1.76 | 1.92 | 2.08 | 2.24 | - |

*Escherichia coli strains, derived from MG1655 with insertions of one out of four TEM variants; Ancestor = TEM-1, Single mutant = G238S, Double mutant = E104K/G238S, Triple mutant = E104K/M182T/G238S.

### S2. Inferring establishment probability from the data.

The single-cell establishment probability is denoted $p_e$. Multiple lineages are assumed to establish independently in the model. The inferred $p_e$ values from multiple inoculum sizes agree, thus validating the assumption (see below).

*Liquid*

The inoculum size is assumed to be Poisson distributed with mean $\lambda$. If the multiple cell lineages grow independently, the establishment probability in a single well is given by

$$p_W = 1 - e^{-\lambda p_e}. \qquad \text{(ES1)}$$

If the total number of wells is $N$, then the mean number of wells with establishment $n = p_W N$; therefore

$$p_e = \frac{-1}{\lambda} \ln(1 - \frac{n}{N}). \qquad \text{(ES2)}$$

We use a bootstrapping algorithm to estimate $p_e$. We do not know $\lambda$ with certainty, so it has to be estimated as well. We generate a total of $R$ resamplings, and each resampling is generated as follows: firstly, a resample is created for the inoculum size by sampling (with replacement) from the CFU (colony forming unit) data for cells dropped on agar, and the mean inoculum size is calculated to obtain a value for $\lambda$; next, a resample (with replacement) is created from the data of the experiment in liquid, and $n$, the number of wells showing establishment, is computed from here. We then use the maximum likelihood estimate of $p_e$ which is

$$\min\{-\frac{1}{\lambda}\ln(1-\frac{n}{N}),1\}. \quad \text{(ES3)}$$

This completes the estimation of $p_e$ from one resample. The final value of $p_e$ is the mean over $R$ resamples created in this way, and the error estimate is the standard deviation of $p_e$ calculated for these $R$ values of $p_e$. We have used $R = 10^5$. The estimates of $p_e$ for low and high inoculum sizes agreed with each other for all mutants and antibiotic concentrations, consistent with our assumption that the cell lineages grow independently (we would have seen a discrepancy between the inferred $p_e$ from low and high inoculum sizes otherwise). The final $p_e$ was obtained as the mean of the $p_e$ for low and high inoculum sizes. The errors of each inoculum size were estimated as the standard deviation of the values of $p_e$ generated through the resampling method. The errors in the final $p_e$ values are the estimated standard deviation of the combined $p_e$. For the double mutant in liquid medium, the highest concentration showed no growth in any well. It was therefore impossible to obtain a non-zero estimate for the error in this case through the bootstrapping method. For curve-fitting, this error was set to be equal to the error estimated at the second-highest concentration.

*Agar*

It is assumed that at 0 concentration, all cells establish colonies. Thus the $p_e(x)$ is estimated simply as $n\,(x)/n\,(0)$ where $n(x)$ is the CFU count at concentration $x$.

### S3. Simple branching process model

To understand the origins of stochasticity, the simplest process one can consider is a branching model where each cell divides independently of the rest of the population with probability $p_d$, and dies with probability $1 - p_d$. Thus the cell population either increases or decreases by 1 at each step. The probability $p_d$ is assumed to be uniform and constant throughout the experiment. The process stops when the population reaches either 0 or a threshold population $n_{\text{th}}$. In the latter case, we say that the population has established. This process is a simple biased random walk starting at population size 1 and exiting through one of two absorbing boundaries. The probability of establishment $p_e$ is the probability that it exits through the boundary at $n_{\mathrm{th}}$, which is

$$p_e = \frac{2-\frac{1}{p_d}}{1-\left(\frac{1}{p_d}-1\right)^{n_{\mathrm{th}}}}. \qquad \text{(ES4)}$$

In the limit $n_{\mathrm{th}} \to \infty$, we obtain the result $p_e = 2 - 1/p_d$ for $p_d \geq \frac{1}{2}$ and 0 otherwise. This limit corresponds to the condition that the population continues to exist indefinitely. This is the relation that has been used in the main text, particularly in Figure 2 (solid symbols with dashed lines). The way $p_d$ approaches $\frac{1}{2}$ asymptotically with increasing concentrations is counter to the expectation that $p_d \to 0$ at very high concentration. To illustrate this point more clearly, we have shown in Figure S1 two possible and qualitatively different scenarios for the relationship between $p_e$ and $p_d$ under the assumptions of our simple branching model. Figure S1a shows the case where $p_e$ approaches $\frac{1}{2}$ asymptotically and $p_e$ goes to zero smoothly. Figure S1b shows the scenario of $p_d$ approaching 0 asymptotically and consequently $p_e$ dropping to zero abruptly. While the latter scenario is the more realistic one, we observe the scenario in Figure S1a. This indicates that one or more of the assumptions in the simple branching model is incorrect. One may reconsider the lack of interaction among cell lineages, but we have shown that interaction between lineages is insignificant in liquid medium. Although the same is not directly demonstrated for agar, the close similarity of the $p_e(x)$ curves for liquid and agar makes this unlikely in the agar medium as well. Another possibility is to adopt a more realistic definition of establishment by requiring that the growth process only needs to reach a certain threshold cell number in the time window of the experiment (Schenk *et al.* 2012). In other words, we take $n_{\mathrm{th}}$ to be finite. We consider the case $n_{\mathrm{th}} = 10^5$, and show in Figure S2 that at this value of $n_{\mathrm{th}}$ the relationship between $p_e$ and $p_d$ is nearly identical to that of the original branching model. This threshold is smaller than the number of cells at which a bacterial population can usually be detected visibly. Choice of a higher value of $n_{\mathrm{th}}$ will make the results even closer to the branching model with infinite threshold. Therefore, considering a finite threshold for establishment does not resolve the issue.

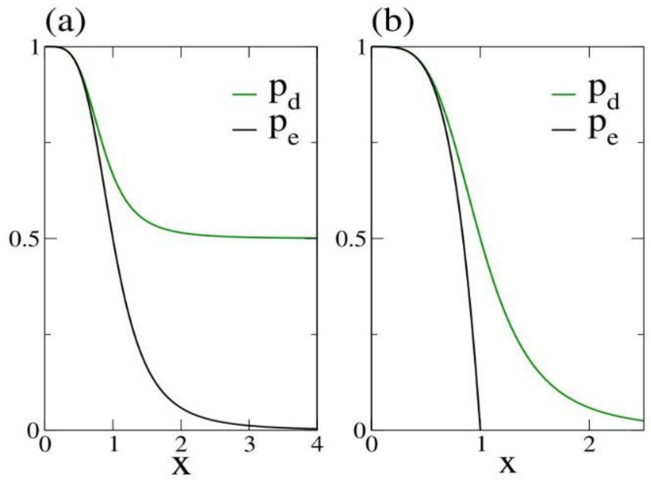


**Figure S1.** Schematic plots showing two different kinds of behavior of $p_e$ (single-cell establishment probability) and $p_d$ (cell division probability) in a simple branching model. x denotes the antibiotic concentration. The two quantities are related by Eq (ES4) with an infinite threshold. **a:** A smooth approach of $p_e$ to zero is linked with $p_d$ approaching the value of $\frac{1}{2}$ asymptotically. This scenario is unrealistic since at very high antibiotic concentration one expects the cell division probability to become zero. **b:** When $p_d$ goes to zero at high concentrations, $p_e$ should become zero abruptly, i.e. at a finite value of x.

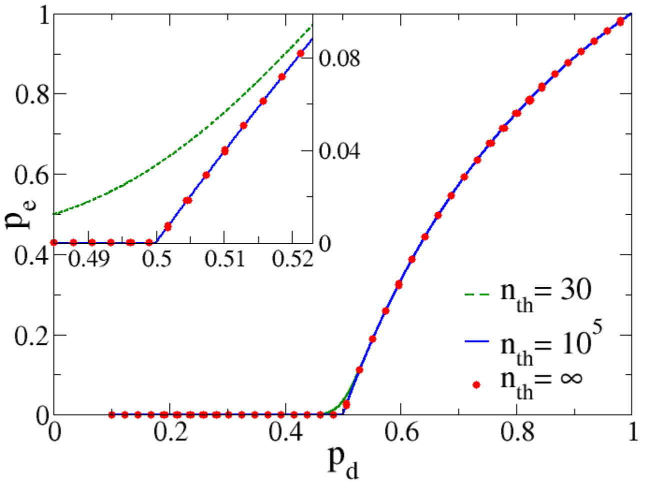


**Figure S2.** $p_e$ as a function of $p_d$ for three different values of the parameter $n_{\text{th}}$. The curves for $n_{\text{th}} = 10^5$ and $n_{\text{th}} = \infty$ (i.e. the original branching model) are nearly identical. The curve for $n_{\text{th}} = 30$ is already close to that of the original branching model. The inset shows a close-up of the graph near the transition point $p_d = 0.5$.

S4. Model with lineage variability

To arrive at a model that incorporates lineage variability, we start by modeling $p_d$ as a Hill function of the antibiotic concentration $x$:

$$p_d(x, x_d) = \frac{1}{1+\left(\frac{x}{x_d}\right)^n}. \qquad \text{(ES5)}$$

Note that in (ES5), $x_d$ is the concentration at which $p_d$ drops to $\frac{1}{2}$, and consequently $p_e$ drops to $0$; in other words, $x_d$ is the single-cell MIC (i.e. the concentration where the growth of a single cell is prevented, as opposed to the standard MIC, which is measured at the population level).

Variability across lineages is introduced via among-lineage variation in the parameter $x_d$. Specifically, we model $x_d$ as a random variable drawn independently for each lineage from the gamma distribution:

$$P(x_d) = \frac{1}{\Gamma(k)a^{k-1}} x_d^{k-1} e^{\frac{-x_d}{a}}.$$

The mean of $x_d$ is $ka$, and the variance is $ka^2$. The parameter $a$ provides a scaling factor of $x_d$. The coefficient of variation of $x_d$ is a measure of lineage variability, which in this case is $k^{\frac{-1}{2}}$; $k$ is therefore inversely related to the degree of lineage variability. The single-cell establishment probability inferred from a large number of observations is then simply the establishment probability (ES5) averaged over the distribution of $x_d$. The final model for the establishment probability is then:

$$p_e(x) = \int_x^\infty dx_d\, P(x_d)\left(1 - \left(\frac{x}{x_d}\right)^n\right). \qquad \text{(ES6)}$$

The parameter $n$ is assumed to be the same for all mutants and both liquid and agar environments. The parameters for the distribution of $x_d$ can be different for the tested strains and can depend on the environment as well. We performed collective fits using the same value of $n$ for all data sets and obtained the estimate $n = 7.18 \pm 0.60$. In Figure S3, the $x$-values for each strain have been scaled by the corresponding estimate for mean $x_d$. The curves for different strains largely collapse, indicating small (but non-zero) lineage variability in $x_d$ within strains. The coefficients of variation of $x_d$, which contain the effect of lineage variability, are larger for the agar than for the liquid environment in most cases, consistent with the less steep curves on agar, but the difference is small.

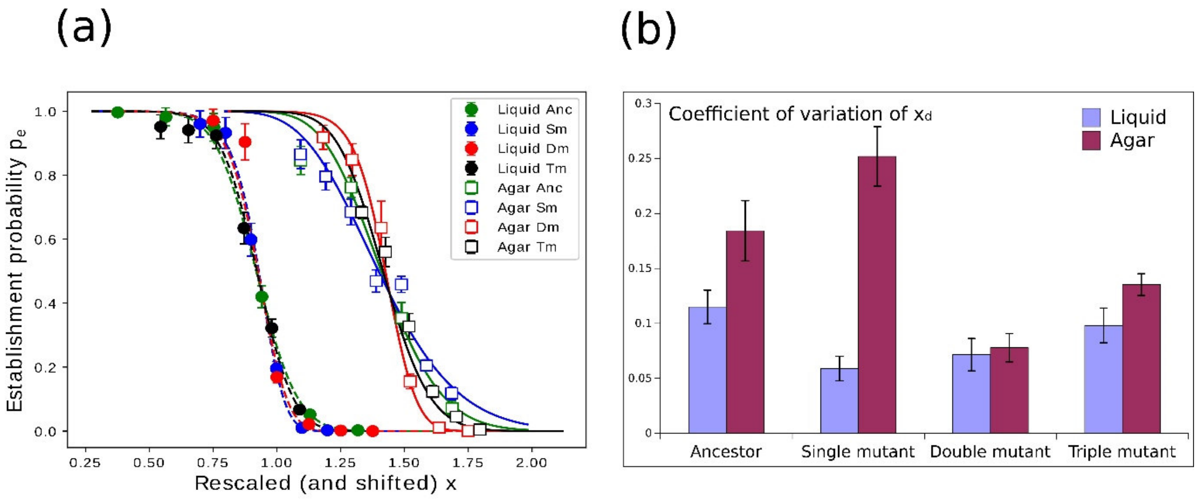


**Figure S3.** Fits with a functional form where cell division probability $p_d$ varies across lineages, and is zero at high concentrations. A joint fit of equation (ES6) with the same value for $n$ for both environments was used. **a**: $p_e(x)$ obtained from curve-fitting. The $x$-values for each strain (Anc: Ancestor, Sm: Single mutant, Dm: Double mutant, Tm: Triple mutant) have been scaled by their respective mean $x_d$, and the values for agar have been shifted along the $x$-axis to the right by 0.5 to improve visibility only. **b**: The coefficient of variation of $x_d$ (mean ± S.D.) measures the degree of lineage variability. It is higher on agar than in liquid for most strains.

S5. Curve fitting for the establishment probability.

The inferred $p_e$, weighted with the estimated errors, were fit by the model described in the main text. The fitting was done using a non-linear least-squares method implemented using the *lmfit* package in python (https://lmfit.github.io/lmfit-py/).

**Table S2:** Parameters of the fits of the establishment probability $p_e(x)$ shown in Figures 2 and 3. $x_0$ is reported in units of the largest concentration used in the liquid and agar experiments for each TEM strain.

| **Strain** | **Experimental environment** | $x_0$ | $n$ | **Pearson correlation of $x_0$ and $n$** |
|---|---|---|---|---|
| Ancestor | liquid | 0.70 +/- 0.006 (0.81%) | 15.33 +/- 0.785 (5.12%) | 0.676 |
| | agar | 0.76 +/- 0.019 (2.54%) | 8.47 +/- 1.311 (15.49%) | <0.001 |
| Single mutant | liquid | 0.77 +/- 0.009 (1.15%) | 23.75 +/- 2.527 (10.64%) | 0.703 |
| | agar | 0.76 +/- 0.018 (2.32%) | 6.24 +/- 0.852 (13.67%) | 0.568 |

| | | | | |
|---|---|---|---|---|
| Double mutant | liquid | 0.68 +/- 0.005 (0.73%) | 22.59 +/- 1.772 (7.84%) | 0.767 |
| | agar | 0.75 +/- 0.013 (1.72%) | 23.57 +/- 2.805 (11.90%) | 0.800 |
| Triple mutant | liquid | 0.84 +/- 0.010 (1.14%) | 14.69 +/- 1.335 (9.09%) | 0.734 |
| | agar | 0.696 +/- 0.014 (1.99%) | 12.891 +/- 1.357 (10.53%) | 0.343 |

S6. Derivation of an expression for $\mathrm{MIC}_q(N)$ for non-interacting lineages.

We define $\mathrm{MIC}_q$ as the concentration at which there is a probability $q$ that no growth will occur. For example, $\mathrm{MIC}_{\frac{2}{3}}$ would be the concentration at which there is $\frac{1}{3}$ probability of growth, and on average 1 out of 3 experiments will show growth. One can then try to estimate $\mathrm{MIC}_q$ for any preferred value of $q$(see Schenk et al., 2012, for the related concept of IC99.99, which was defined as the concentration that killed 99.99% of the population). The MIC further depends on the inoculum size $N$ (Brook 1989). Importantly, the basic inoculum effect does not require social interactions but is also apparent when the cell division probability $p_d$ is independent of density. Social interactions, as expected for β-lactamases such as TEM, can amplify the inoculum effect, for which it is important to have a "null expectation" of the inoculum effect in the absence of social effects (Alexander and MacLean 2020). Therefore, a more informative quantity to look at it is $\mathrm{MIC}_q(N)$, i.e. the antibiotic concentration where the probability of no growth is equal to $q$ for an inoculum of $N$ cells. A population fails to establish if none of the $N$ cells establishes, and assuming that the cell lineages do not interact, the probability that a population does not establish is the product of the probabilities that each cell in the inoculum does not establish.

For the establishment probability, we have used the fitting function

$$p_e = \frac{1}{1+\left(\frac{x}{x_0}\right)^n}, \qquad \text{(ES7)}$$

where each strain has its own $x_0$ and $n$. Now we define $\mathrm{MIC}_q(N)$ as the antibiotic concentration $x$ at which the probability that an inoculum of $N$ cells fails to establish is $q$. When cell lineages do not interact, the probability that none of $N$ cells establishes is $(1-p_e)^N$. Then, $\mathrm{MIC}_q(N)$ is the solution $x$ of

$$\left(\frac{\left(\frac{x}{x_0}\right)^n}{1+\left(\frac{x}{x_0}\right)^n}\right)^N = q.$$

Therefore,

$$\mathrm{MIC}_q(N) = x_0 \left( \frac{q^{\frac{1}{N}}}{1-q^{\frac{1}{N}}} \right)^{\frac{1}{n}}. \qquad \text{(ES8)}$$

For large $N$ (and provided $q$ is not equal to 0 or 1), one can make the asymptotic approximation:

$$\frac{\mathrm{MIC}_q(N)}{x_0} \cong \left( \frac{N}{\ln\frac{1}{q}} \right)^{\frac{1}{n}}, \qquad \text{(ES9)}$$

and therefore, to leading order,

$$\ln \frac{\mathrm{MIC}_q(N)}{x_0} \cong \frac{1}{n} \ln N$$

The null-model predictions for $\mathrm{MIC}_q(N)$ in the main text were obtained using the exact equation (ES8). Notice from the last equation that the leading term in $\ln\frac{\mathrm{MIC}_q(N)}{x_0}$ does not depend on $q$. However, this approximation is valid only when $q$ is not close to 0 or 1.

The probability that no growth is observed at concentration $x$ for an inoculum of size $N$, assuming independence of lineages, is $\left(1 - p_e(x)\right)^N$; therefore the likelihood that growth is observed at $x$ is $1 - \left(1 - p_e(x)\right)^N$. The MIC assays were performed in duplicate, and the probability that growth is observed for a particular $x$ in both is

$$L = \left(1 - \left(1 - p_e(x)\right)^N\right)^2.$$

This formula was used for the null-model likelihood $L$ in Table 1 in the main text. The fitted curves of $p_e(x)$ were used in the formula. The value of $x$ used was the highest concentration at which growth was observed in the MIC assay (i.e. half the determined MIC due to the 2-fold CTX dilution).

### References


Alexander H. K., and R. C. MacLean, 2020 Stochastic bacterial population dynamics restrict the establishment of antibiotic resistance from single cells. Proc. Natl. Acad. Sci. U. S. A. 117: 19455–19464. https://doi.org/10.1073/pnas.1919672117

Brook I., 1989 Inoculum Effect. Rev. Infect. Dis. 11: 361–368. https://doi.org/10.1093/clinids/11.3.361

Schenk M. F., I. G. Szendro, J. Krug, and J. A. G. M. de Visser, 2012 Quantifying the adaptive potential of an antibiotic resistance enzyme. PLoS Genet. 8: e1002783. https://doi.org/10.1371/journal.pgen.1002783
